# GABAergic Circuit Activation Induces a Therapeutically Responsive State for anti-PD-L1 Immunotherapy in Glioblastoma

**DOI:** 10.64898/2026.08.26.747200

**Authors:** Marta Scalera, Elisa De Santis, Filippo Rossi, Nicolò Meneghetti, Lorenzo Amir Nemati Fard, Pasquale Miglionico, Francesco Raimondi, Alessandra Flori, Massimo Pasqualetti, Luca Menichetti, Soma Sengupta, Eleonora Vannini, Mario Costa

**Author notes:** Corresponding authors: Eleonora Vannini, Mario Costa. Last author: Mario Costa.

## Abstract

Glioblastoma (GBM) disrupts cortical excitatory-inhibitory balance and establishes an immunosuppressive microenvironment that limits therapeutic efficacy. Whether restoring inhibitory signaling can restrain glioma progression and improve responsiveness to immune checkpoint blockade remains unknown. Peritumoral parvalbumin-positive (PV^+^) interneurons were bidirectionally manipulated by chemogenetics in orthotopic GL261 gliomas to assess tumor growth and neurological function. GABA_B_ signaling was pharmacologically activated with baclofen in GL261 and CT-2A models and combined with anti-PD-L1 blockade in GL261. Therapeutic response, survival, tumor rechallenge, and early myeloid remodeling were evaluated. Human GBM single-cell transcriptomic data were analyzed to examine the relationship between GABAergic and PD-L1 intercellular signaling. PV activation transiently restrained glioma growth, reduced tumor proliferation and preserved cortical function, whereas PV^+^ silencing increased seizure susceptibility and neurological impairment without accelerating tumor growth. Baclofen monotherapy did not affect survival, whereas its combination with anti-PD-L1 immunotherapy induced complete tumor eradication in 66% of GL261-bearing mice, prolonged survival, and conferred durable protection against tumor rechallenge. Combination therapy also altered the proportions of Arg1+ and CD11c+ cells within the intratumoral F4/80+ compartment. Human single-cell analysis revealed a shared myeloid-centered communication axis linking GABA_B_ and PD-L1 signaling. These findings identify GABAergic signaling as a modulator of GBM progression and demonstrate that combining baclofen with anti-PD-L1 induces durable tumor regression, and prolongs survival in the GL261 model, supporting a neuro-immune framework for combining GABAergic modulation with immunotherapy.

## Introduction

Glioblastoma (GBM) remains the most lethal and biologically aggressive primary brain tumor in adults, characterized by diffuse infiltration, profound therapeutic resistance, and a median survival of less than 15 months despite maximal treatment ^1^. Although therapeutic strategies have traditionally focused on tumor-intrinsic oncogenic mechanisms, it is increasingly recognized that GBM progression is critically shaped by reciprocal interactions with the surrounding neural and immune microenvironments ^2^. Malignant glioma cells establish functional synaptic and paracrine communication with host neurons, integrating into neural circuits through activity-dependent and AMPA receptor-mediated mechanisms that promote tumor growth, invasion, and network remodeling ^3,4^.

A key consequence of tumor–neural crosstalk is the development of peritumoral network hyperexcitability. Glioma cells increase extracellular glutamate through enhanced release and impaired astrocytic uptake, creating an excitotoxic environment that supports tumor progression while contributing to neurological dysfunction, including tumor-associated epilepsy ^5^. Although the pro-tumoral role of excitatory glutamatergic signaling is well established, the contribution of inhibitory neuronal circuits to GBM biology remains less defined.

Parvalbumin-positive (PV^+^) interneurons are central regulators of cortical excitation– inhibition balance and network synchronization ^6^. In glioma, progressive dysfunction of these fast-spiking interneurons in the peritumoral cortex impairs inhibitory control, increases network excitability, and contributes to neurological dysfunction ^7,8^. Conversely, selective activation of PV^+^ interneurons has been shown to suppress glioma proliferation, suggesting that restoration of inhibitory signaling may represent a therapeutically actionable strategy ^9,10^.

Beyond their effects on tumor growth and network function, neuronal circuits may also influence the immune state of the tumor microenvironment. This possibility is particularly relevant in GBM, where tumor-associated microglia, macrophages, and infiltrating myeloid populations constitute a major component of the tumor microenvironment and contribute to immunosuppression and resistance to immunotherapy ^11^. The glioma-associated myeloid compartment is highly plastic and can adopt distinct functional states in response to local microenvironmental cues. Whether alterations in neuronal inhibitory signaling contribute to this immune regulation, however, remains largely unexplored ^12^.

GABAergic signaling represents a potential molecular interface between neural activity and immune regulation. GABA receptors are expressed across multiple cellular compartments of the tumor microenvironment, and emerging evidence indicates that GABAergic pathways can modulate immune-cell function ^13,14^. These observations raise the possibility that restoring inhibitory signaling may influence GBM not only through neuronal and tumor-associated mechanisms, but also by altering the immune microenvironment. Baclofen, a clinically established GABA_B_ receptor agonist, provides a pharmacological strategy to test this hypothesis and to determine whether GABAergic modulation can enhance responsiveness to immune checkpoint blockade.

Here, we investigated whether strengthening peritumoral inhibitory signaling can restrain glioma progression, preserve neurological function, and enhance antitumor immunity. We combined cell-specific chemogenetic manipulation of PV^+^ interneurons with pharmacological activation of GABA_B_ receptors in immunocompetent GBM models to define the contribution of inhibitory signaling to tumor progression and neurological dysfunction. We then tested whether GABA_B_ receptor activation could enhance the therapeutic efficacy of PD-L1 blockade and examined the associated remodeling of the tumor immune microenvironment. Finally, we analyzed human GBM single-cell data to determine whether GABAergic and immune checkpoint signaling are linked within shared intercellular communication networks. Together, our findings identify inhibitory signaling as a neuroimmune regulator of GBM and provide a preclinical rationale for combining GABAergic modulation with immune checkpoint blockade.

## Materials and Methods

### Animal Models and Ethical Statements

Adult (age > postnatal day 60) wild type male and female C57BL/6 J and B6.129P2-*Pvalb^tm^*^1^*^(cre)Arbr^*/J mice were bred in our animal facility and housed in a 12 h light/dark cycle, with food and water available ad libitum. All experimental procedures respected the ARRIVE guidelines and the European Communities Council Directive #86/609/EEC and were approved by the Italian Ministry of Health (981/2020-PR, released on 11/03/2016 and 433/2026-PR, released on June 16, 2026). Male and female animals were equally distributed across all experimental groups.

### Stereotaxic Adeno-Associated Virus (AAV) Injections

B6.129P2-Pvalbtm1(cre)Arbr/J mice were anesthetized with an intraperitoneal (ip) injection of a ketamine (100 mg/kg) and xylazine (10 mg/kg) and placed on a stereotaxic apparatus. To selectively manipulate PV^+^ interneurons, mice received intracranial injections of Cre-dependent AAV vectors encoding mCherry-tagged DREADDs. PV^+^ activation was achieved using AAV8-hSyn-DIO-hM3D(Gq)-mCherry (Addgene #44361-AAV8; titer ≥ 4×10¹² vg/mL), whereas AAV9-hSyn-DIO-hM4D(Gi)-mCherry (Addgene #44362-AAV9; titer ≥ 1×10¹³ vg/mL) was administered for inhibition.

Viral delivery was performed using a Nanoliter 2020 Injector (World Precision Instruments) at a constant infusion rate of 90 nL/min. To achieve widespread expression within the motor cortex, injections were made at two mediolateral coordinates (1200 μm and 2300 μm lateral to Bregma) and at two depths per site (600 μm and 900 μm below the dural surface). A volume of 300 nL was delivered at each depth, yielding a total of 1200 nL. The injection needle was left in place for 5 min after each infusion before slow withdrawal to minimize reflux. Glioma cells were implanted two weeks after AAV injection.

### Glioma cell culture

Murine glioma GL261 and CT-2A cell lines were cultured as monolayers in Dulbecco’s Modified Eagle Medium (DMEM) F-12 supplemented with 10% fetal bovine serum (FBS), 1% penicillin/streptomycin, and 1% L-glutamine. Cells were maintained at 37°C in a humidified atmosphere containing 5% CO2. The culture medium was refreshed three times per week, and cells were passaged upon reaching 80%–90% confluency. All cell lines were routinely tested for mycoplasma contamination and used within 10 passages from thawing.

### Orthotopic glioma inoculation

Syngeneic tumor induction was performed as previously described in ^9^. Mice were anesthetized ip with ketamine (100 mg/kg) and xylazine (10 mg/kg) and positioned in a stereotaxic frame. A total of 40,000 GL261 cells (20,000 cells/μL in PBS) or 50,000 CT-2A cells (25,000 cells/μL in PBS) were stereotaxically implanted into either the primary motor cortex (1.75 mm lateral and 0.5 mm anterior to Bregma) or the primary visual cortex (2.5 mm lateral to the midline, at the level of Lambda). A total volume of 2 μL of cell suspension was slowly delivered at a depth of 0.9 mm below the pial surface using a Hamilton syringe connected to an automated infusion pump (KD Scientific) at a constant rate of 0.2 μL/min. Following injection, the needle was left in place for 5 minutes before slow withdrawal to prevent tumor cell reflux. The implantation site and cell line were selected according to the specific experimental paradigm.

### Chemogenetic treatments and CNO administration

For the *in vivo* chemogenetic manipulation of PV^+^ interneurons, Clozapine N-oxide dihydrochloride (CNO; Tocris Bioscience, cat. no. #6329) was dissolved in sterile 0.9% saline and administered ip at 1 mg/kg. Control animals received an equivalent volume of vehicle (0.9% saline). CNO/saline was administered twice daily (morning and evening) from day 12 to day 22 after tumor implantation.

### BrdU administration and tissue collection

To label proliferating cells, mice received a single ip injection of 5-bromo-2’-deoxyuridine (BrdU; Sigma-Aldrich) at 50 mg/kg on day 22 after glioma implantation. Twenty-four hours later, mice were deeply anesthetized, transcardially perfused and brains were collected and processed for immunofluorescence analyses.

### Immune Checkpoint Inhibition

Orthotopic glioma-bearing mice received ip injections of an anti-PD-L1 antibody (200 μg/mouse; Cat. #BE0101, Bio X Cell) twice weekly from day 12 to day 28 after tumor induction.

### Immunofluorescence

Glioma-bearing mice were deeply anesthetized and transcardially perfused with phosphate-buffered saline (PBS; Sigma-Aldrich), followed by 4% paraformaldehyde (PFA) in 0.1 M sodium phosphate buffer (pH 7.4). Brains were carefully removed, post-fixed in the same fixative for 4 h at 4 °C, and cryoprotected in a 30% sucrose solution. Coronal sections (50 μm thick) were subsequently obtained using a sliding microtome (Leica).

To validate chemogenetic manipulation, free-floating sections were immunostained with rabbit anti-Parvalbumin (Cat# 195 004, 1:500; Synaptic Systems) and rabbit anti-c-Fos (Cat. # 226 008, 1:1000; Synaptic Systems) primary antibodies, followed by Alexa Fluor 488-conjugated anti-rabbit secondary antibody. Endogenous mCherry fluorescence was visualized in coronal sections without signal amplification.

To assess GBM cell proliferation, brains were collected 23 days after tumor implantation, following BrdU administration. For antigen retrieval, free-floating sections were incubated in 2 M HCl for 30 min at 37 °C and subsequently washed three times in PBS for 10 min each. Sections were then incubated with rat anti-BrdU primary antibody (Cat. #ab6326, 1:500; Abcam), followed by an Alexa Fluor 488-conjugated anti-rat secondary antibody (Cat# A-11006, 1:1000; Invitrogen).

To characterize the tumor-associated myeloid compartment, free-floating sections underwent antigen retrieval in Tris-EDTA buffer for 5 min at 90 °C and were immunostained with rat anti-F4/80 (Cat. #ab6640, 1:500; Abcam) and rabbit anti-Arginase-1 (Cat. #PA5-95956, 1:100; Invitrogen) or rabbit anti-CD11c (Cat. #HS-375 003, 1:100; SySy). F4/80 was detected using a Cy3-conjugated anti-rat secondary antibody (1:500), whereas Arg1 or CD11c with Alexa Fluor 488-conjugated anti-rabbit secondary antibody. Cell nuclei were counterstained with Hoechst (1:500; Bisbenzimide, Sigma-Aldrich), and sections were mounted with Vectashield (Vector Laboratories). All experimental groups were processed and acquired under identical staining and imaging conditions.

### Fluorescence Microscopy and Quantitative Image Analysis

DREADD expression and BrdU incorporation were acquired using a Zeiss Axio Observer microscope equipped with an AxioCam MRm camera (Carl Zeiss MicroImaging GmbH), using 10× and 40× objectives, respectively. Myeloid markers were acquired using a Zeiss LSM 900 confocal microscope equipped with an Airyscan 2 detector and a 40× oil-immersion objective. For all experimental groups, image acquisition parameters were kept identical, including laser power, detector gain, and acquisition settings.

To evaluate the specificity of viral transduction, PV^+^ interneurons and PV/mCherry double-positive cells were manually quantified within predefined regions of interest (ROIs) using the Cell Counter plugin in ImageJ (National Institutes of Health). Transduction efficiency was calculated for each animal as the percentage of PV^+^/mCherry^+^ cells relative to the total PV^+^ population. Tumor cell proliferation was assessed by quantifying the proportion of BrdU^+^ cells within the tumor area. The proliferation index was calculated as the ratio of BrdU^+^ cells to the total number of Hoechst-positive nuclei within selected ROIs, as previously described in ^15^. Myeloid cells were quantified using an automated QuPath workflow to identify F4/80^+^/Arg1^+^ and F4/80^+^/CD11c^+^ cells within tumor ROIs. At least six animals per group were analyzed. Measurements from multiple sections and ROIs were averaged for each animal, which was considered one biological replicate for statistical analysis.

### *In Vitro* Clonogenic Assay

GL261 and CT-2A glioma cells were detached by trypsinization, resuspended in complete culture medium, and counted using a Bürker hemocytometer (Marienfeld). Cells were seeded into 12-well plates at a density of 300 cells per well and allowed to adhere overnight. The following day, cultures were treated with increasing concentrations (50–200 μM) of baclofen ((±)-β-(aminomethyl)-4-chlorobenzenepropanoic acid; Sigma-Aldrich, Cat. #B5399] or γ-aminobutyric acid (GABA; Sigma-Aldrich) and maintained under standard culture conditions for 9 days. At the end of the treatment period, colonies were fixed and stained with crystal violet. Colonies containing at least 50 cells were manually counted under a light microscope to assess, and clonogenic survival was expressed as the number of colonies relative to untreated control.

### Primary Cortical Astrocyte Isolation and Culture

Primary cortical astrocytes were isolated from postnatal day 1–3 (P1–P3) mouse pups as previously described in (Streubel-Gallasch et al., 2021). Briefly, brains were collected in ice-cold Dulbecco’s phosphate-buffered saline (DPBS; Biowest), and the olfactory bulbs, meninges were removed. Cerebral cortices were mechanically dissociated by passing them through a 70 μm cell strainer (Sarstedt) using a syringe plunger in Dulbecco’s modified Eagle medium (DMEM; Sigma-Aldrich) supplemented with 10% fetal bovine serum (FBS) and 1% penicillin/streptomycin.

Cell suspensions were centrifuged at 300 x g for 15 min, washed twice with 25 mL of complete culture medium and seeded into T75 culture flasks. Culture medium was replenished after 7 days and subsequently every 3–4 days. Once cultures reached approximately 80% confluence, microglia were removed by orbital shaking, and the remaining astrocyte monolayer was harvested for downstream experiments. All experiments were performed using at least three independent primary astrocyte cultures derived from separate litters.

### RNA Extraction and Quantitative Real-Time PCR (qRT-PCR)

Total RNA was extracted from cultured cells using the PureLink RNA Mini Kit (Invitrogen; Thermo Fisher Scientific) according to the manufacturer’s instructions. RNA concentration and purity were determined using a NanoDrop 2000 spectrophotometer (Thermo Fisher Scientific).

Complementary DNA (cDNA) was synthesized from equal amounts of total RNA using the High-Capacity cDNA Reverse Transcription Kit (Applied Biosystems; Thermo Fisher Scientific). Quantitative real-time PCR (qRT-PCR) was performed using the PowerUp SYBR Green Master Mix on a 7500 Fast Real-Time PCR System (Applied Biosystems). Relative gene expression was calculated using the 2^-ΔΔCt^ method with β-actin as the endogenous reference gene. Primer sequences used for qRT-PCR are provided in **Supplementary Table 1**.

### *In vivo* baclofen administration

Baclofen (Sigma-Aldrich, Cat# B5399) was dissolved in 1 M HCl to prepare a stock solution, which was diluted in sterile saline immediately before use. Mice received daily ip injections of baclofen (1 mg/kg) or an equivalent volume of vehicle. Treatment was administered from day 12 to day 28 post-tumor implantation in the GL261 model and from day 7 to day 28 in the CT-2A model.

### Motor Assessments

Motor performance was evaluated using the grip strength and the grid walk tests, as previously described in ^16,17^. Baseline performance was assessed 15 days prior to tumor inoculation. Motor testing was performed twice weekly, starting on day 10 for GL261-bearing mice and day 7 for CT-2A-bearing mice. All assessments were conducted at the same time of day by investigators blinded to treatment allocation. Longitudinal measurements were normalized to each animal’s baseline performance.

### Grip Strength Assay

Forelimb grip strength was measured using a digital Grip Strength Meter (GSM; SKU No. 47200, Ugo Basile S.R.L). Mice were allowed to grasp the grid with their forepaws while being gently pulled backward by the tail until grip release. The maximal force exerted before release was automatically recorded. Three consecutive trials were performed for each animal at each time point, and the average value was used for analysis.

### Grid Walk Assay

Fine motor coordination was assessed using an elevated wire grid (32 cm × 20 cm with 11 mm × 11 mm mesh). Mice were allowed to freely explore the apparatus for 5 min, and sessions were video-recorded. Foot slips were quantified offline in a blinded manner and scored separately for the limbs contralateral and ipsilateral to the tumor. The percentage of foot slips in the affected limb was calculated as the number of foot-slip errors divided by the total number of steps taken by the affected limb, multiplied by 100.

### Post-treatment behavioral monitoring

Following each intraperitoneal injection of CNO or vehicle, mice were visually observed for 30 min, starting 30 min after treatment, for the occurrence of generalized tonic–clonic seizures. Seizures were identified based on overt bilateral convulsive movements accompanied by loss of postural control, based on established behavioral criteria ^18^.

### Electrophysiological Recordings and Visually Evoked Potentials (VEPs)

In vivo electrophysiological recordings were performed exclusively in the experimental cohort undergoing chemogenetic activation of PV^+^ interneurons, to evaluate visual cortical function during GBM progression. Vehicle- and clozapine-N-oxide (CNO)-treated mice were anesthetized with ketamine (100 mg/kg) and xylazine (10 mg/kg, i.p.) and placed in a stereotaxic frame. Following a craniotomy over the tumor-bearing visual cortex (3.6–4.1 mm lateral to lambda), a 16-channel linear silicon probe (NeuroNexus) was inserted into the cortex for local field potential (LFP) recordings. Neural signals were acquired using an OmniPlex recording system (Plexon) and analyzed with NeuroExplorer software. Visual stimuli were generated using PsychoPy and presented on a gamma-corrected monitor to stimulate the binocular visual field. Phase-reversing horizontal square-wave gratings (1 Hz) of varying spatial frequencies (0.06–0.5 cycles/degree) and contrasts (10–90%) were presented as previously described ^19–22^. VEP amplitude and latency were quantified from the principal negative (N1) and positive (P1) components. Visual acuity and contrast sensitivity were determined by linear extrapolation of VEP amplitude as a function of spatial frequency and stimulus contrast, respectively.

### *In Vivo* Magnetic Resonance Imaging (MRI)

Tumor growth was longitudinally monitored using a 7-T vertical bore preclinical MRI scanner (Avance III HD 300 MHz; Bruker BioSpin) equipped with a 30-mm ^1H quadrature volume coil. Mice were anesthetized with ketamine/xylazine (100/10 mg/kg, i.p.), and body temperature and respiration were continuously monitored throughout image acquisition. MRI scans were performed without contrast agent on days 12, 22, and 28 after GL261 implantation and on days 7, 14, 21, and 28 after CT-2A implantation. The imaging protocol included FLASH scout images for anatomical localization and T2-weighted TurboRARE sequences for tumor visualization and volumetric analysis. Tumor boundaries were manually segmented on consecutive MRI slices using Mango software (version 4.1), and total tumor volume was calculated by integrating the segmented areas across slices and expressed in mm^3^.

### Single-cell Data Collection and Processing

The Core GBMap dataset ^23^ was retrieved from CELLxGENE. Expression was denoised and batch-corrected using scvi-tools (v1.4.0) in Python (v3.12.11) ^24^. An scVI model (n_layers = 2, n_latent = 30, gene_likelihood = “nb”) was trained on the raw counts using all genes and default parameters, treating each sample as a separate batch. Normalized expression was obtained with the get_normalized_expression method (library size set to 10,000) and log1p-transformed. To batch-correct the data, we used scVI’s counterfactual prediction via the transform_batch argument. To avoid decoding cells out of the training distribution (i.e. conditioning a cell on a batch in which few similar cells were observed), we processed each cell type separately and considered as valid only those batches containing at least 5 cells of that type. Each cell’s normalized expression was thus computed as the mean of its decoded expression across all batches in which at least 5 cells of the same type were present.

### Cell-Cell Communication Inference

The denoised expression was used to infer cell-cell communication with the CellPhoneDB ^25^ method implemented in the LIANA (v1.6.1) Python package ^26^, using the ligand-receptor reference from ^27^ (which aggregates interactions from CellPhoneDB ^25^, CellChat ^28^, and the GPCR Cancer Axes resource ^29^. Because CellPhoneDB records GABBR1 and GABBR2 interactions as separate entries (e.g. GAD1+SLC32A1-GABBR1 and GAD1+SLC32A1-GABBR2), individual GABBR1 and GABBR2 entries were merged into a single GABBR1+GABBR2 complex. Significance was assessed using Benjamini-Hochberg-corrected p-values (p < 0.05), and non-significant interactions were discarded. For each source-target cell-type pair, all ligand–receptor pairs involving the GABBR1+GABBR2 complex were summed into a single GABAergic signaling score; analogously, a PD-L1 signaling score was obtained by summing all interactions with CD274 as their ligand.

The CellPhoneDB analysis was then repeated separately for each sample. In this per-sample analysis, expression was again denoised with get_normalized_expression, but each cell was decoded conditioning on its own original batch, so as not to mask between-sample heterogeneity. A separate score (CellPhoneDB’s lr_means value) was obtained for each combination of source, target, ligand, and receptor in each sample; non-significant interactions (Benjamini-Hochberg-corrected p < 0.05) were set to zero. Principal component analysis (scikit-learn, v1.7.2) was performed on the resulting interaction × sample matrix, with samples as observations and interactions as features and without additional standardization, identifying 44 principal components that cumulatively explained 90% of the variance in the cell-cell communication networks. The total strength of GABA_B_ and PD-L1 signaling was computed for each sample by summing all interactions involving the GABBR1+GABBR2 complex and the CD274 gene, respectively. Spearman correlation coefficients between each principal component and the two signaling scores were computed (scipy.stats.spearmanr, SciPy v1.18.0), and associations significant after Benjamini-Hochberg correction across all tested component–pathway pairs (p < 0.05) were retained to identify components jointly associated with both pathways.

### Statistical Analysis

Statistical analyses were performed using GraphPad Prism 8.0 (GraphPad Software Inc.). Data distribution was assessed for normality before parametric analyses. Comparisons between two groups were performed using unpaired two-tailed Student’s t-test, whereas comparisons involving three or more groups were analyzed by one-way ANOVA followed by Tukey’s multiple-comparison test. Longitudinal motor and imaging data were analyzed using two-way repeated-measures ANOVA, followed by the Holm–Sidak multiple-comparison test. Survival was analyzed using Kaplan–Meier curves and compared with the long-rank (Mantel-Cox) test. Data are presented ± SEM unless otherwise indicated. In all analyses, a P-value < 0.05 was considered statistically significant.

## Results

### Peritumoral PV^+^ interneuron stimulation transiently restrains glioma growth and mitigates cortical dysfunction

To investigate whether activating peritumoral PV^+^ interneurons influences GBM progression, PV-Cre mice received injections of AAV-hSyn-DIO-hM3D(Gq)-mCherry into the motor cortex surrounding the tumor implantation site. Histological analyses confirmed robust and selective DREADD expression in PV^+^ interneurons, with approximately 80% of PV^+^ cells co-expressing mCherry (**Fig. 1A, B**; *P* < .001). Furthermore, systemic administration of clozapine N-oxide (CNO) markedly increased c-Fos expression in hM3D(Gq)-expressing PV interneurons, confirming effective chemogenetic activation (**Fig. 1C**).

**Figure 1:**
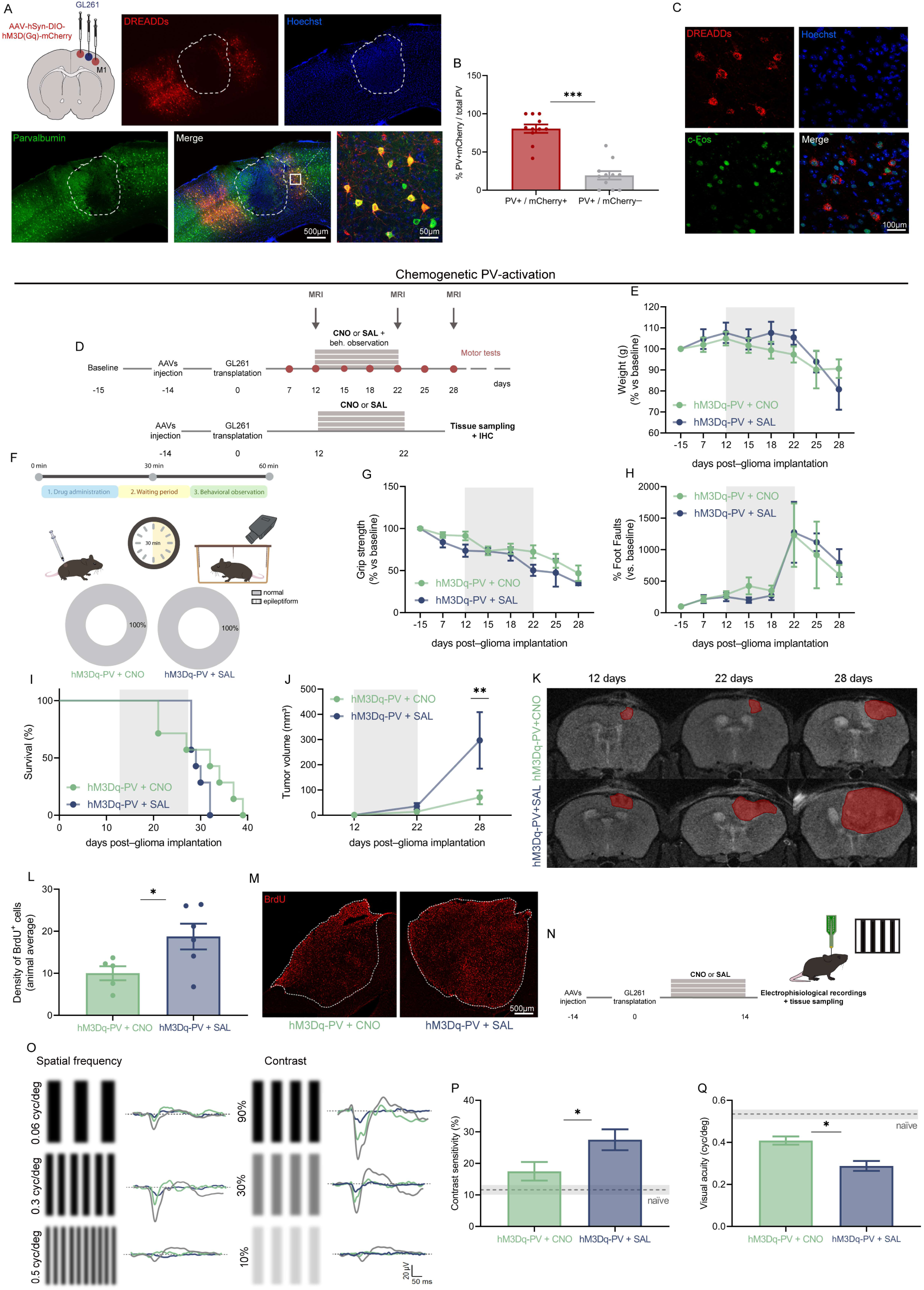
Transient activation of peritumoral PV^+^ interneurons limits glioma progression and preserves cortical function. (**A**) Schematic of bilateral AAV-hSyn-DIO-hM3D(Gq)-mCherry microinjections into the motor cortex surrounding the glioma site. Representative immunofluorescence images showing selective expression of the excitatory DREADD (mCherry, red) within PV^+^ interneurons (green); nuclei are counterstained with Hoechst (blue). Dotted lines outline the tumor core. (**B**) Quantitative analysis of viral targeting efficiency, confirming that approximately 80% of peritumoral PV^+^ interneurons express mCherry (*P* < .001, Student’s t-test). (**C**) Representative immunohistochemical validation of functional activation showing robust c-Fos expression (green) selectively colocalizing with mCherry-expressing PV^+^ interneurons (red) after clozapine N-oxide (CNO) administration. (**D**) Experimental timelines. The upper panel details the protocol for chemogenetic manipulation alongside survival, MRI, and motor function readouts; the lower panel outlines the experimental protocol for immunohistochemical cell proliferation analyses. (**E**) Longitudinal monitoring of body weight, showing stable and comparable profiles between the hM3Dq-PV + CNO (n = 7) and hM3Dq-PV + SAL (n = 7) groups. (**F**) Behavioral monitoring protocol (top) and quantification of seizure activity (bottom); 100% of mice in both cohorts exhibit normal behavior with no detectable epileptiform activity. (**G**) Grip strength and (**H**) grid walk (% foot faults) performance across tumor progression, showing no significant differences between treatment arms. Data in E, G, and H represent mean ± SEM, analyzed by two-way repeated-measures ANOVA, normalized to baseline (day -15). (**I**) Kaplan–Meier survival curves indicating no significant divergence between hM3Dq-PV + SAL (n = 7) and hM3Dq-PV + CNO (n = 7) glioma-bearing mice (log-rank Mantel–Cox test). (**J**) Volumetric quantification of longitudinal tumor growth expressed in mm^3^. Chemogenetic stimulation yields a significant reduction in tumor burden at day 28 in the hM3Dq-PV + CNO cohort compared with vehicle controls (*P* = .004, two-way repeated-measures ANOVA followed by Tukey’s post-hoc test). Values are normalized to day 12. (**K**) Representative serial coronal T2-weighted MRI scans of a PV-activated and a vehicle-treated mouse across timepoints (days 12, 22, and 28). Red overlays outline the segmented tumor mass. (**L**) Quantitative density of BrdU+ proliferating cells within the tumor mass at day 23 post-injection, showing a significant decrease in the PV-activated group (*P* = .043, Student’s t-test). (**M**) Representative overview images of intratumoral BrdU incorporation (red) in hM3Dq-PV + CNO and hM3Dq-PV + SAL mice. Dotted lines define tumor boundaries. (**N**) Experimental timeline for in vivo extracellular electrophysiological recordings in the primary visual cortex at day 15 post-injection. (**O**) Graphical representation of visual stimuli (varying spatial frequencies and contrasts) and corresponding representative visual evoked potential (VEP) traces; green lines represent VEPs from PV-activated mice (hM3Dq-PV + CNO), blue lines represent vehicle-treated controls (hM3Dq-PV + SAL), and gray lines represent tumor-free, non-injected naïve mice. (**P**) Quantitative analysis of contrast sensitivity, demonstrating significant preservation of visual processing capacity in PV-activated mice compared to vehicle-treated controls (*P* < .05, two-way repeated-measures ANOVA followed by Holm– Sidak post-hoc test; horizontal dashed line indicates naïve baseline). (**Q**) Volumetric-matched visual acuity quantification, showing preserved spatial frequency discrimination in the hM3Dq-PV + CNO cohort compared to the vehicle group (*P* < .05, two-way repeated-measures ANOVA followed by Holm–Sidak post-hoc test; solid horizontal line indicates naïve baseline).

Following MRI confirmation of tumor engraftment at day 12, GL261-bearing mice were randomized to receive daily CNO (1 mg/kg, i.p) or saline (SAL) from day 12 to day 22 post-tumor induction (Vannini et al. 2017; **Fig. 1D**). Chemogenetic activation of peritumoral PV^+^ interneurons did not adversely affect the general health of the animals. Physiological parameters, including body weight and motor performance, remained stable and comparable to the hM3Dq + SAL control group (**Fig. 1E, G, H**). Moreover, visual monitoring of freely moving animals after CNO administration revealed no generalized tonic-clonic seizures throughout the treatment period (**Fig. 1F**).

Longitudinal MRI analysis revealed that activation of peritumoral PV^+^ interneurons markedly delayed GBM progression. Whereas tumors in saline-treated mice expanded progressively, tumor growth remained largely stable throughout the treatment period in CNO-treated animals, resulting in a significantly lower tumor volume at day 28 (**Fig. 1J, K**; *P* = .004). However, rapid tumor progression occurred following treatment discontinuation, indicating that tumor growth suppression was not sustained after cessation of PV+ interneuron activation.

To determine whether reduced tumor growth was associated with decreased tumor cell proliferation, BrdU incorporation was quantified at the end of the treatment period. Chemogenetic activation significantly reduced the density of BrdU+ cells within the tumor compared with saline-treated controls (*P* = .043; **Fig. 1L, M**), indicating that PV+ interneuron activation reduced glioma cell proliferation. Despite these effects, PV^+^ interneuron activation did not significantly prolong overall survival (**Fig. 1I**).

Overall, these results indicate that increasing the activity of peritumoral PV^+^ interneurons transiently attenuates glioma progression and reduces tumor cell proliferation, but is insufficient to achieve durable control of tumor growth.

We next evaluated whether preserving inhibitory circuit activity could also mitigate tumor-induced cortical dysfunction. GL261 cells were implanted into the primary visual cortex, and visually evoked potentials (VEPs) were recorded from the peritumoral cortex 15 days after tumor implantation (**Fig. 1N**). A cohort of naïve, tumor-free mice was included as physiological controls. All experimental groups—hM3Dq-PV + CNO, hM3Dq-PV + SAL, and naïve mice—were presented with visual stimuli varying in contrast (10% to 90%) and spatial frequency (0.06 to 0.5 cycles/degree; **Fig. 1O**) to quantify contrast sensitivity and visual acuity. Although visual function remained impaired relative to naïve animals, mice receiving CNO displayed significantly greater visual acuity and contrast sensitivity than saline-treated glioma-bearing mice (**Fig. 1P, Q**). These findings indicate that activation of peritumoral PV^+^ interneurons not only transiently limits glioma growth but also partially preserves cortical sensory processing during tumor progression.

### Chemogenetic silencing of peritumoral PV^+^ interneurons exacerbates neurological dysfunction without altering GBM progression

To complement the activation studies, we next investigated the effects of chemogenetic silencing of peritumoral PV^+^ interneurons during glioma progression. PV-Cre mice received bilateral injections of AAV-hSyn-DIO-hM4D(Gi)-mCherry and underwent the same experimental timeline and analyses as the activation cohort (**Fig. 2A**). To exclude potential off-target effects of CNO, an additional cohort of tumor-bearing mice lacking DREADD expression received either CNO or saline. No differences in body weight, motor performance, survival or tumor growth were observed between these groups (**Supplementary Fig. 1**).

**Figure 2.**
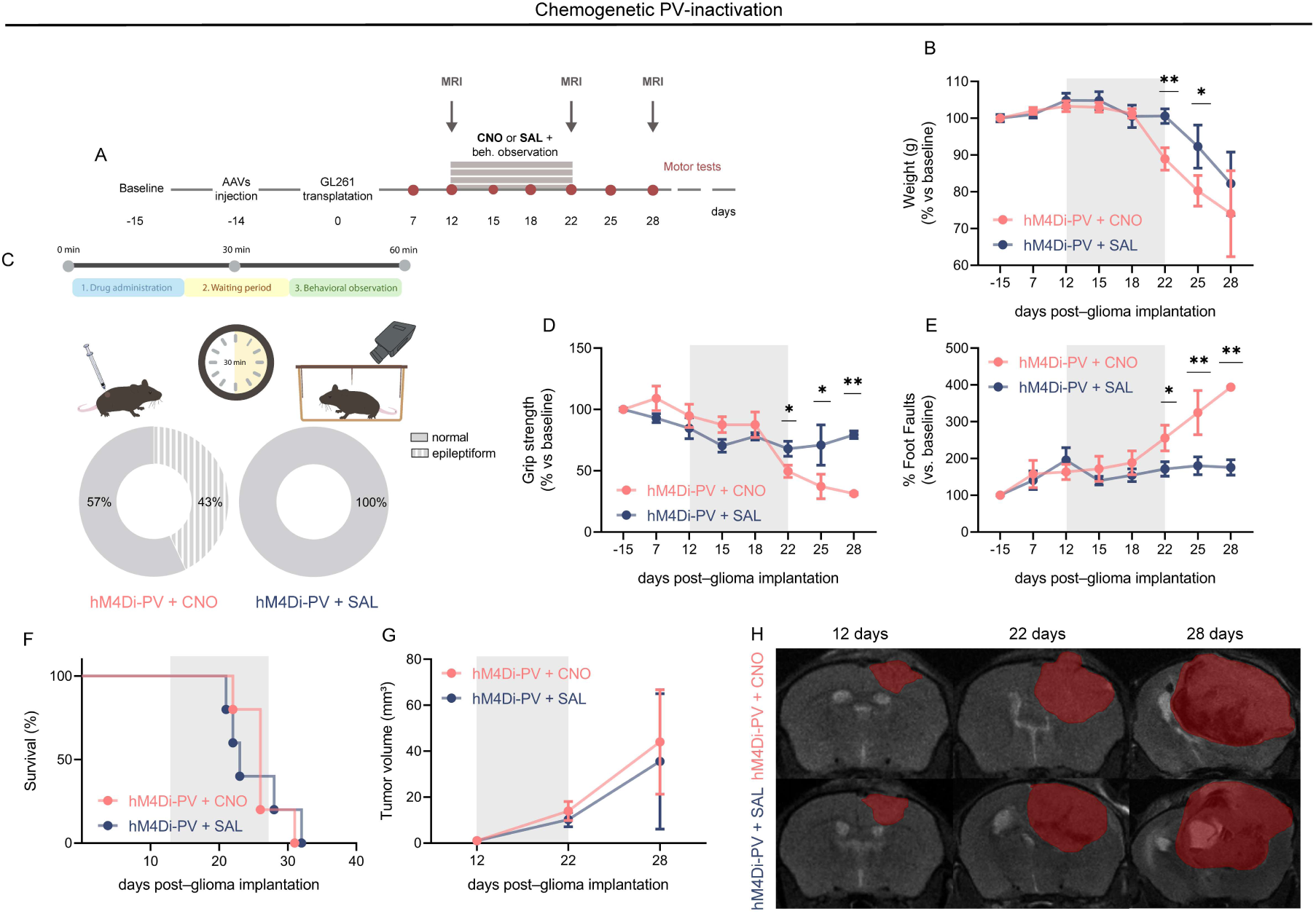
Chemogenetic inactivation of peritumoral PV-interneurons exacerbates GBM-associated neurological deficits without affecting tumor progression. **(A)** Experimental timeline. (**B**) Longitudinal monitoring of body weight, showing an accelerated weight loss in PV-inactivated glioma-bearing mice (hM4Di-PV + CNO, n = 7) at day 22 (*P* = .002) and day 25 (*P* = .010) compared with vehicle controls (hM4Di-PV + saline, n = 6). (**C**) Behavioral monitoring protocol (top) and quantification of seizure activity (bottom); 43% (n = 3/7) of mice in the hM4Di-PV + CNO cohort exhibit behavioral epileptiform activity post-CNO administration, while 100% of vehicle-treated controls (hM4Di-PV + SAL) display normal behavior. (**D**) Grip strength and (**E**) grid walk (% foot faults) performance across tumor progression. PV-inactivated mice display significant motor impairment at 22 and 25 days post-glioma implantation compared with vehicle controls in both the grip strength test (D; *P* = .034 at 22 days; *P* = .029 at 25 days) and the grid walk test (E; *P* = .048 at 22 days; *P* = .001 at 25 days). Data in B, D, and E represent means ± SEM, analyzed by two-way repeated-measures ANOVA, normalized to baseline (day -15). (**F**) Kaplan–Meier survival curves indicating no significant divergence between hM4Di-PV + SAL (n = 6) and hM4Di-PV + CNO (n = 7) glioma-bearing mice (log-rank Mantel–Cox test, *P* = .464). (**G**) Volumetric quantification of longitudinal tumor growth expressed in cubic millimeters (mm^3^). Chemogenetic inhibition yields no significant differences in tumor burden kinetics between the vehicle and PV-inactivated cohorts (two-way repeated-measures ANOVA). Values are normalized to day 12. (**H**) Representative serial coronal T2-weighted MRI scans of a PV-inactivated mouse (top row) and a vehicle control mouse (bottom row) across timepoints (days 12, 22, and 28). Red overlays outline the segmented tumor mass.

Chemogenetic silencing of PV^+^ interneurons resulted in a pronounced worsening of the neurological phenotype. Compared with saline-treated controls, hM4Di-PV mice receiving CNO exhibited significantly greater body weight loss (**Fig. 2B**; *P* = .002 at day 22; *P* = .010 at day 25), progressive impairment of forelimb grip strength (**Fig. 2D**; *P* = 0.034 at day 22; *P* = 0.029 at day 25; *P* = 0.010 at day 28) and increased motor coordination deficits in the grid walk test (**Fig. 2E**; *P* = .048 at day 22; *P* = .001 at day 25; *P* = .010 at day 28). Notably, behavioral observation during the 30-min period following CNO administration identified generalized tonic-clonic seizures in 43% of PV-silenced mice, whereas no seizures were observed in control animals (**Fig. 2C**), indicating that endogenous PV^+^ interneuron activity contributes to limiting tumor-associated neurological dysfunction.

In striking contrast to the severe neurological deterioration, chemogenetic silencing of PV^+^ interneurons did not alter GBM progression. Longitudinal MRI showed comparable tumor growth between CNO- and saline-treated mice throughout the experimental period (**Fig. 2G, H**), and no differences in overall survival were observed (**Fig. 2F**). Taken together, these findings reveal a functional dissociation between neurological deterioration and tumor progression. While PV^+^ interneuron silencing did not significantly alter GBM progression, it markedly exacerbated neurological dysfunction, indicating that the effects of PV activity on tumor progression and cortical function can be dissociated.

### Systemic baclofen treatment exerts model-dependent anti-tumor activity

To translate our chemogenetic findings into a clinically relevant therapeutic strategy, we investigated systemic activation of GABAergic signaling using baclofen, an FDA-approved GABA_B_ receptor agonist widely used for the treatment of spasticity ^31^.

We first examined whether baclofen could directly target GBM cells. Quantitative RT-PCR analysis showed robust expression of GABA_B_ receptor subunit GABA_B_R1 (Gabbr1) in primary murine astrocytes as well as in GL261 and CT-2A glioma cells (**Fig. 3A-C**). In contrast, GABA_B_R2 (Gabbr2), which is required for canonical GABA_B_ receptor signaling, was detected in primary astrocytes but was undetectable in either GBM cell lines. Consistent with these findings, treatment with increasing concentrations of baclofen or GABA (50-200 μM) failed to affect clonogenic growth of either GL261 or CT-2A cells (**Fig. 3D-H**), indicating that baclofen does not directly impair GBM cell proliferation *in vitro*.

**Figure 3.**
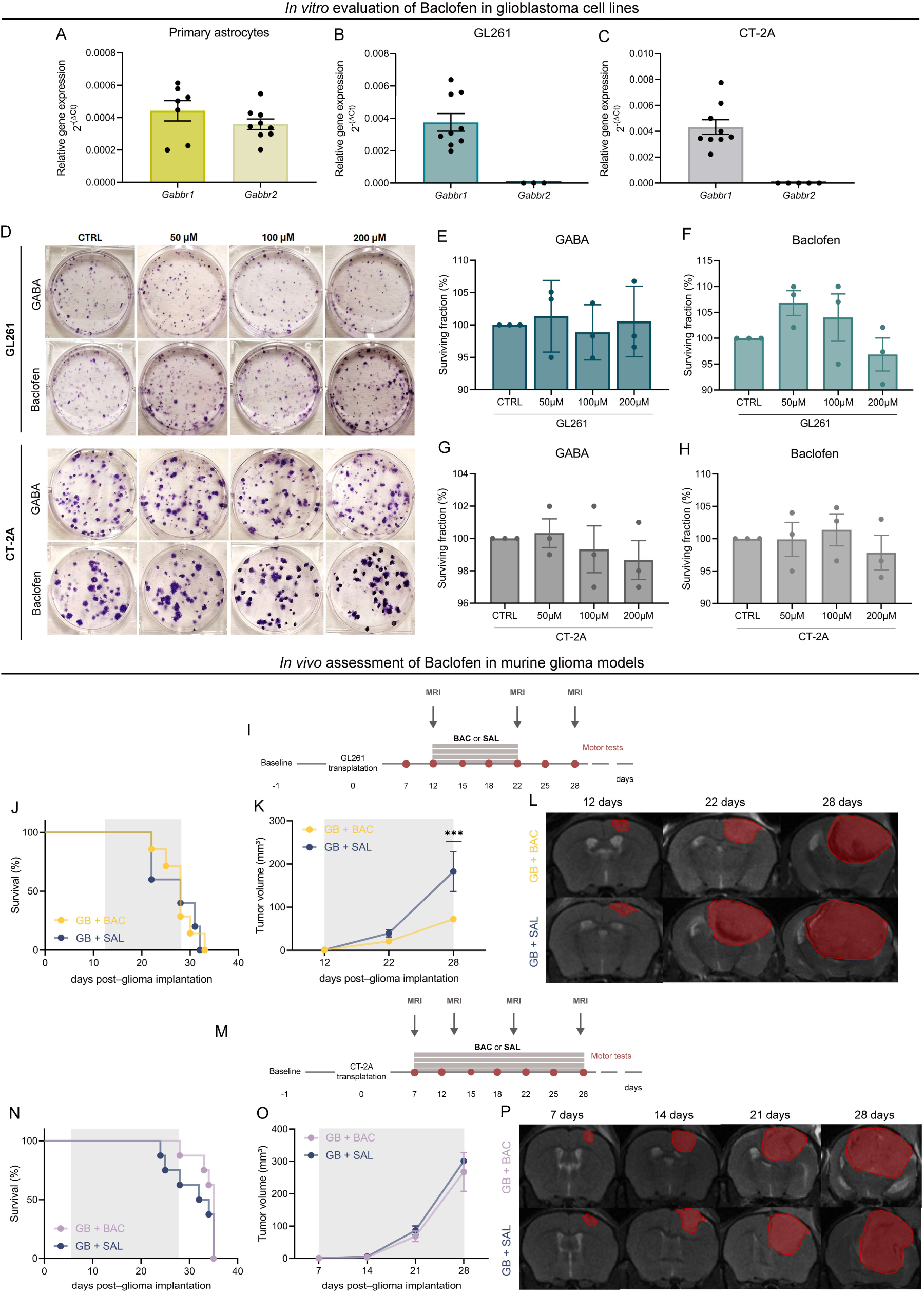
Baclofen treatment reduces tumor growth in the GL261 model without affecting glioma cell viability in vitro. (**A–C**) Relative gene expression levels (2-^ΔCt^) of Gabbr1 and Gabbr2 in primary astrocytes (**A**), GL261 cells (**B**), and CT-2A cells (**C**), as determined by qRT-PCR. While Gabbr1 is robustly expressed across all evaluated cell types, Gabbr2 transcripts are detected in primary astrocytes but remain completely undetectable in both murine glioma cell lines. Data are presented as mean ± SEM of three independent biological replicates; statistical significance was evaluated using Student’s t-test on ΔCT values. (**D**) Representative images of crystal violet-stained colony formation assays under baseline (CTRL) conditions or challenging cells with GABA or baclofen at designated concentrations (50 μM, 100 μM, and 200 μM) in GL261 and CT-2A lines. (**E–H)** Quantitative assessment of the surviving fraction (%) in GL261 cells exposed to GABA (**E**) or baclofen (**F**), and CT-2A cells treated with GABA (**G**) or baclofen (**H**). No statistically significant variations in colony numbers are observed compared with untreated control groups (Student’s t-test). Data represent mean ± SEM of three technical replicates per condition. (**I**) Experimental timeline illustrating the experimental paradigm for GL261 glioma-bearing mice subjected to longitudinal MRI, vehicle (SAL), or baclofen (BAC) pharmacotherapy and motor assessment. (**J**) Kaplan–Meier survival curves indicating no significant survival differences between baclofen-treated mice (GB + BAC, yellow line, n = 8) and vehicle controls (GB + SAL, blue line, n = 6) within the GL261 model (log-rank Mantel–Cox test). (**K**) MRI-based volumetric quantification of longitudinal GL261 tumor progression; the vehicle cohort displays a significantly elevated tumor volume compared with the baclofen-treated group at the final experimental timepoint (day 28 post-implantation; *P* < .004, two-way repeated-measures ANOVA). Values are normalized to the first imaging session at day 12. (**L**) Representative serial coronal T2-weighted MR images of GL261-bearing mice from both experimental arms across day 12, 22, and 28 timepoints. Red masks highlight the manually segmented tumor boundaries. (**M**) Schematic representation of the in vivo experimental design and treatment schedule optimized for the CT-2A syngeneic model. (**N**) Kaplan–Meier survival curves for CT-2A glioma-bearing mice, revealing overlapping survival distribution profiles between the GB + BAC (violet line, n = 8) and GB + SAL (blue line, n = 8) cohorts. (**O**) Volumetric growth kinetics derived from longitudinal MRI data for the CT-2A cohort, demonstrating no significant deviations in tumor progression between vehicle-treated and baclofen-treated groups at any analyzed timepoint (two-way repeated-measures ANOVA). Data are normalized to day 7 baseline imaging. (**P**) Representative serial coronal T2-weighted MRI scans tracking CT-2A tumor expansion at days 7, 14, 21, and 28 post-injection, with red overlays delineating the progressive space-occupying lesion.

Given the absence of a detectable direct effect *in vitro*, we next investigated whether systemic GABA_B_ receptor activation could influence GBM progression *in vivo*. Baclofen was administered daily using an extended treatment schedule (days 12-28 after tumor implantation in the GL261 model and days 7-28 in the CT-2A model; **Fig. 3I, 3M**). Although baclofen modestly attenuated neurological decline, improving grid walk performance in GL261 mice (*P* = .0010 at day 28) and forelimb grip strength in CT-2A mice (*P* = .0379 at day 24; **Supplementary Fig. 2**), its anti-tumor efficacy differed substantially between models.

Longitudinal MRI revealed a significant reduction in tumor volume in baclofen-treated GL261 mice at the final imaging time point (*P* = .004; **Fig. 3K, 3L**), whereas no effect on tumor burden was detected in the more aggressive CT-2A model (**Fig. 3O, 3P**). Consistent with these imaging findings, baclofen treatment failed to prolong overall survival in either GBM models (**Fig. 3J, 3N**).

Together, these findings argue against a direct antitumor effect of baclofen on GL261 or CT-2A cells *in vitro* and suggest that its *in vivo* activity may involve indirect effects mediated by the tumor microenvironment. However, the magnitude of this benefit is highly context-dependent and, as monotherapy, remains insufficient to achieve durable tumor control.

### Combined baclofen and PD-L1 blockade induces durable tumor regression and long-term immunological memory

Given the modest therapeutic benefit of baclofen monotherapy, we next investigated the therapeutic response to combined GABA_B_ receptor activation and PD-L1 blockade. GL261-bearing mice received vehicle, anti-PD-L1 monotherapy, or baclofen plus anti-PD-L1(**Fig. 4A, Supplementary Figure 3A**).

**Figure 4.**
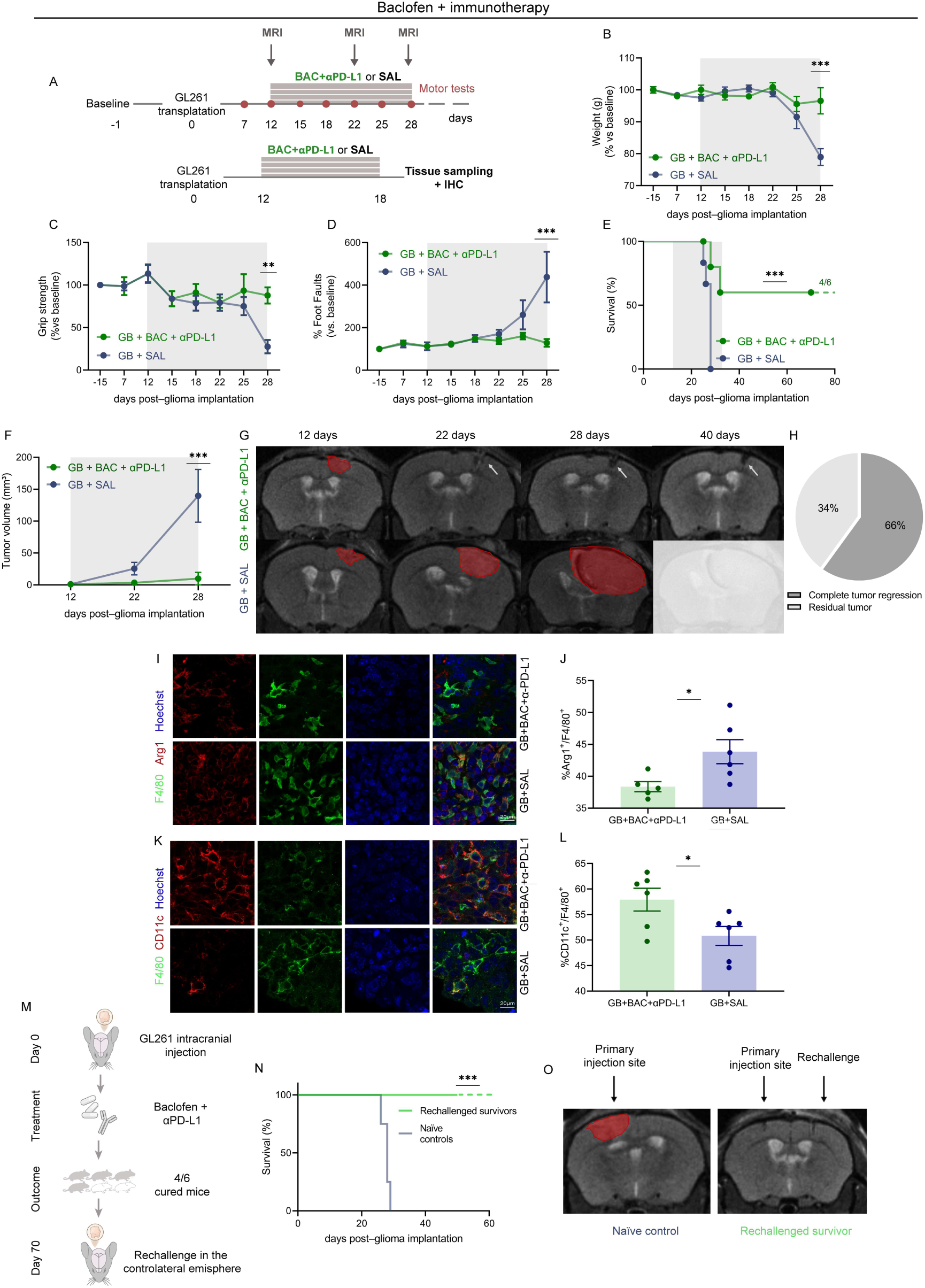
Combined baclofen and PD-L1 blockade induces durable tumor regression and protection against tumor rechallenge with early changes in the F4/80+ myeloid compartment. (**A**) Experimental design. GL261-bearing mice received baclofen (BAC; 1 mg/kg, i.p.) daily and anti-PD-L1 antibody twice weekly from day 12 to day 28 after tumor implantation. A separate cohort was treated from day 12 to day 18 for immunohistochemical analyses of the tumor immune microenvironment. SAL, saline (vehicle). (**B**) Longitudinal monitoring of body weight changes, expressed as a percentage of baseline values, demonstrating stable physiological conditions in mice treated with baclofen plus anti-PD-L1 (GB + BAC + αPD-L1, green line, n = 6) compared with vehicle-treated controls (GB + SAL, blue line, n = 6; *P* < .001). (**C**) Grip strength and (**D**) Grid walk (% foot faults) performance during tumor progression, demonstrating preserved motor function in the combination therapy group compared with the progressive neurological decline observed in vehicle-treated mice (*P* < .01 for grip strength; *P* < .001 for grid walk at day 28). Data in B–D are presented as mean ± SEM and were analyzed using two-way repeated-measures ANOVA after normalization to baseline values. (**E**) Kaplan–Meier survival curves showing significantly prolonged overall survival in mice treated with baclofen plus anti-PD-L1 (GB + BAC + αPD-L1, n = 6) compared with vehicle-treated controls (GB + SAL, n = 6; *P* = .0058, log-rank Mantel–Cox test). (**F**) Longitudinal MRI quantification of tumor volume (mm³), demonstrating marked suppression of tumor growth following combination therapy (*P* < .001, two-way repeated-measures ANOVA). Tumor volumes were normalized to day 12. (**G**) Representative serial coronal T2-weighted MR images acquired at days 12, 22, 28, and 40 after glioma implantation in mice treated with baclofen plus anti-PD-L1 (top) or vehicle (bottom). Red overlays indicate segmented tumor boundaries. White arrows indicate complete radiological regression of the tumor. (**H**) Pie chart summarizing tumor response in the combination treatment cohort at day 40, showing complete tumor regression in 66% of treated mice and residual tumor burden in the remaining 34%. (**I**) Representative immunofluorescence images showing F4/80 (green), Arginase-1 (Arg1, red), and Hoechst (blue) staining in tumor sections from vehicle- and baclofen plus anti-PD-L1-treated mice. Scale bar, 20 μm. (**J**) Percentage of Arg1^+^ cells among F4/80^+^ myeloid cells, showing a reduced proportion following combination therapy (*P* = .0344). (**K**) Representative immunofluorescence images showing F4/80 (green), CD11c (red), and Hoechst (blue) staining in tumor sections from vehicle- and baclofen plus anti-PD-L1-treated mice. Scale bar, 20 μm. (**L**) Percentage of CD11c^+^ cells among F4/80^+^ myeloid cells, showing a reduced proportion following combination therapy (*P* = .0334). Quantitative data in J and L are presented as mean ± SEM and were analyzed using an unpaired two-tailed Student’s *t* test. (**M**) Schematic representation of the tumor rechallenge protocol. Mice exhibiting complete tumor regression after combination therapy (4 of 6 mice) received a contralateral intracranial injection of GL261 cells 70 days after the initial tumor implantation without further treatment. (**N**) Kaplan–Meier survival curves following tumor rechallenge, showing complete protection of previously cured mice (green line, n = 4) compared with age-matched naïve controls (blue line, n = 4; *P* < .001, log-rank Mantel–Cox test). (**O**) Representative T2-weighted MR images showing the primary tumor at the initial implantation site (left, red overlay) and the absence of tumor at both the primary and contralateral rechallenge sites following secondary GL261 implantation (right), confirming durable immunological protection.

Combination therapy markedly improved disease course. Treated mice maintained body weight and motor performance, with neurological function progressively recovering toward baseline levels, whereas vehicle-treated animals showed progressive deterioration (**Fig. 4B-D**). Consistent with these functional improvements, combined Baclofen and PD-L1 blockade significantly prolonged overall survival compared with vehicle-treated controls (**Fig. 4E**; *P* = .0058). Longitudinal MRI further revealed a profound therapeutic response to the combined treatment (**Fig. 4F, G**). A complete radiological regression occurred in 66% of treated mice (n = 4/6) and was sustained throughout the 70-day follow-up. In these animals, MRI showed only a residual hypointense cavity at the original tumor site, consistent with tissue remodeling after tumor clearance (**Fig. 4H**). By contrast, anti-PD-L1 monotherapy resulted in long-term survival in 14% of animals (n = 1/7; **Supplementary Figure 3D-E**).

To investigate early immune changes preceding tumor regression, immunohistochemical analysis of the intratumoral F4/80^+^ myeloid compartment was performed (**Fig. 4A**). This analysis revealed a reduced proportion of F4/80^+^ cells expressing Arg1 (**Fig. 4I, J**; *P* = .0344) and CD11c (**Fig. 4K, L**; *P* = .0344) in glioma-bearing mice receiving combination therapy.

Given the durable therapeutic response, we next asked whether complete radiological regression following combination therapy was associated with protective anti-tumor immunity. Mice showing complete radiological regression (4/6) were therefore rechallenged with GL261 cells in the contralateral hemisphere 70 days after the initial tumor implantation (**Fig. 4M**). Age-matched naïve mice receiving a primary tumor implantation served as controls. Strikingly, all four previously responding mice remained tumor-free following rechallenge, whereas age-matched naïve controls developed rapidly growing tumors and exhibited the expected median survival of 28 days (**Fig. 4N**). Consistent with these findings, MRI of rechallenged survivors revealed only the injection tract without evidence of tumor formation (**Fig. 4O**). Together, these data identify the remodeling of the glioma myeloid compartment as a hallmark of the durable therapeutic response elicited by combined Baclofen and PD-L1 blockade.

### A shared myeloid axis links GABAergic and PD-L1 signaling in human GBM single-cell

Given the early myeloid remodeling observed following combined baclofen and PD-L1 blockade, we next investigated whether GABA_B_ and PD-L1 signaling could also be identified within the human GBM microenvironment. We analyzed a publicly available single-cell atlas ^23^ comprising 338,564 cells from 110 patients spanning 17 major cell populations (**Fig. 5A**). GABA_B_ and PD-L1 pathway components showed distinct cell type-specific expression patterns, with PDCD1 largely restricted to T and NK cells and GABBR1/2 enriched in neurons and astrocytes, whereas ligand expression was more broadly distributed across cell populations (**Fig. 5B**). CellPhoneDB analysis identified 1,462 significant interactions involving GABA_B_ or PD-L1 signaling. GABAergic interactions predominantly involved neural populations, whereas PD-L1 signaling was primarily directed toward T and NK cells (**Fig. 5C**). GABA_B_ signaling predominantly targeted neurons, astrocytes and oligodendrocyte precursor cells (238 significant interactions each), whereas neurons (188 interactions), oligodendrocyte precursor cells (125) and malignant cells (114) represented the principal sources of GABAergic ligands. In contrast, PD-L1 signaling was almost exclusively directed toward T cells (17 interactions) and, to a lesser extent, NK cells (16 interactions), while PD-L1 ligands were broadly distributed across multiple cellular compartments. Accordingly, the global communication network was dominated by GABA_B_ interactions converging on neuronal populations and PD-L1 interactions converging on T cells (**Fig. 5C**).

**Figure 5.**
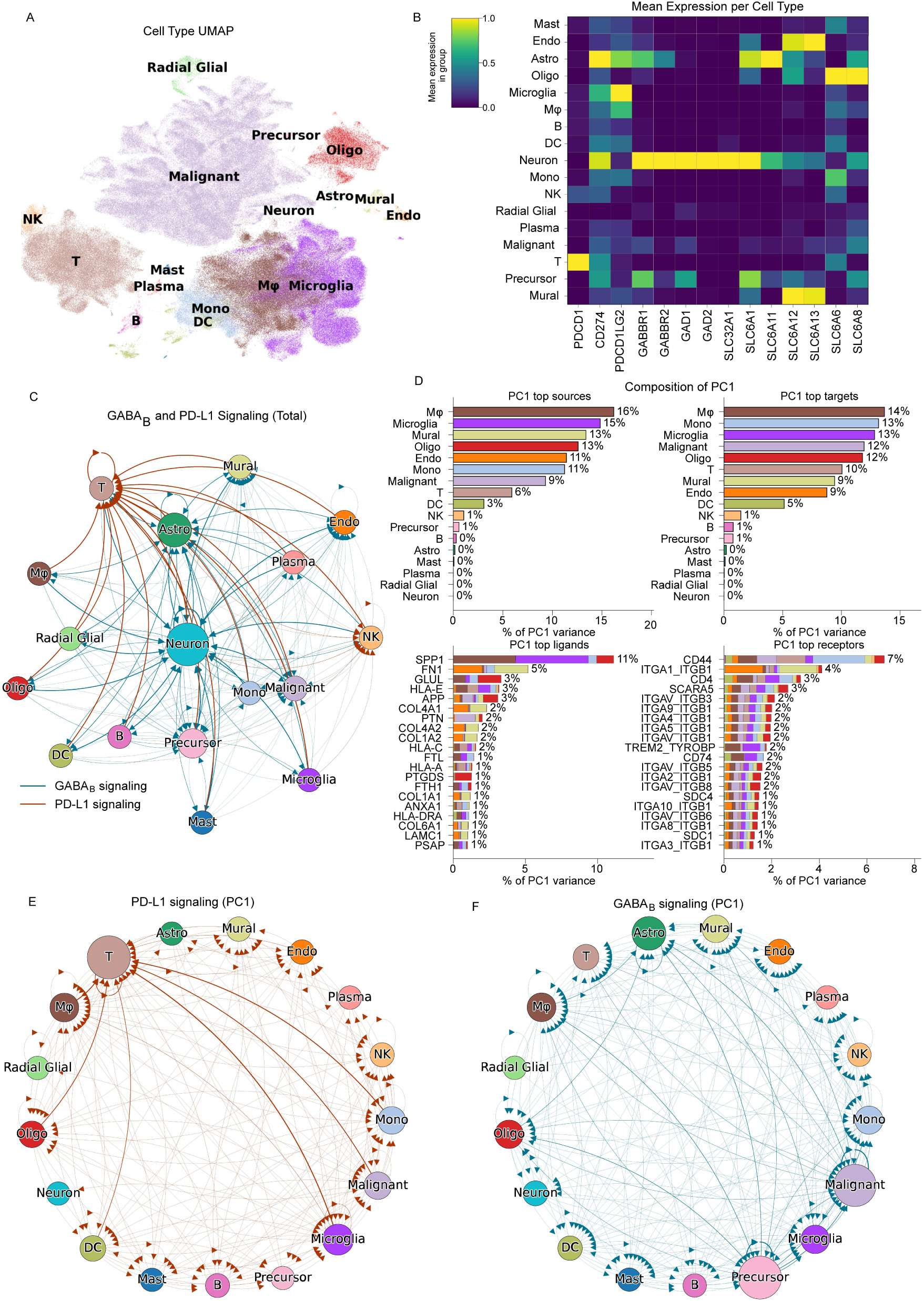
Cell-Cell communication analysis of human GBM single-cell. (**A**) UMAP representation of the human GBM single-cell dataset comprising 338,564 cells from 110 patients across 17 major cell populations, colored by cell type. (**B**) Mean batch-corrected expression of genes involved in GABA_B_ and PD-L1 signaling across cell types. Expression values for each gene are normalized between 0 and 1. PDCD1 expression is predominantly restricted to T and NK cells, whereas GABBR1/2 are enriched in neural populations. (**C**) Network visualization of integrated GABA_B_ (blue) and PD-L1 (red) cell-cell communication. CellPhoneDB identified 1,462 significant ligand-receptor interactions involving the two pathways. GABA_B_ signaling predominantly involves neural populations, whereas PD-L1 signaling is primarily directed toward T and NK cells. Edge weights represent the total interaction strength summed across ligand-receptor pairs associated with each pathway. (**D**) Composition of the first principal component (PC1) of patient-specific communication networks according to source and target cell types, ligands, and receptors. PC1 accounts for 22.6% of inter-sample variance and is the only component significantly associated with both GABA_B_ (Spearman’s ρ = 0.86, *P* = 2.5 × 10⁻³⁹) and PD-L1 signaling (ρ = 0.76, *P* = 9.7 × 10⁻²⁷; Benjamini-Hochberg corrected). The top 20 interactions by contribution to PC1 are shown. Stacked bars indicate the source and target cell-type composition of ligand and receptor contributions, respectively. Interactions contributing to PC1 are predominantly associated with macrophages, microglia, and monocytes and include SPP1-CD44, APP-CD74, MHC-I, integrin-matrix, and TREM2-TYROBP signaling. (**E,F**) Network visualization of PD-L1 (**E**) and GABA_B_ (**F**) components contributing to PC1. PD-L1 interactions predominantly originate from myeloid and tumor-associated populations and converge on T cells, whereas GABAergic interactions connect neural/tumor populations with multiple cellular compartments, including myeloid populations. Edge weights are proportional to the contribution of each interaction to PC1. In all network visualizations, edge width is proportional to edge weight and node size to the sum of incoming and outgoing edge weights.

To determine whether these pathways co-varied across individual tumors, we reconstructed patient-specific cell-cell communication networks and performed principal component analysis. Forty-four principal components accounted for 90% of inter-sample variance. PC1, explaining 22.6% of the variance, was the only component significantly associated with both GABA_B_ (Spearman’s ρ = 0.86, *P* = 2.5 × 10⁻³⁹) and PD-L1 signaling (ρ = 0.76, *P* = 9.7 × 10⁻²⁷; BH corrected), whereas the remaining components were selectively associated with either pathway (**Fig. 5D**). Thus, GABA_B_ and PD-L1 signaling co-varied across human GBMs along a shared axis of intercellular communication. Interactions contributing most strongly to PC1 revealed a myeloid-centered communication network dominated by macrophages, microglia, and monocytes and included SPP1-CD44, APP-CD74, MHC-I, integrin-matrix, and TREM2-TYROBP signaling. Within this shared axis, myeloid populations represented major sources of PD-L1 signaling toward T cells and were also embedded within the GABAergic communication network (**Fig. 5E, 5F**). Together, these findings identify a shared myeloid-centered communication axis along which GABA_B_ and PD-L1 signaling co-vary in human GBM, independently supporting an association between GABAergic and immune signaling within the tumor microenvironment.

## Discussion

GBM remains one of the most aggressive primary brain tumors, with limited therapeutic options and limited clinical benefit from immune checkpoint blockade, despite the success of these therapies in other malignancies ^32–34^. Increasing evidence suggests that this resistance reflects not only profound immune suppression but also the complex integration of neuronal, immune, and tumor-derived signals within the GBM microenvironment ^35,36^.

Here, we identify GABAergic signaling as a potential interface between these compartments. Activation of peritumoral PV+ interneurons restrained glioma progression, while pharmacological activation of GABA_B_ receptors markedly potentiated the efficacy of PD-L1 blockade in the GL261 model. Although these represent distinct experimental manipulations, together they implicate inhibitory GABAergic signaling as a modifiable component of the GBM neuroimmune landscape. Importantly, our data do not establish that the antitumor effects of PV activation are specifically mediated through GABAB receptors. Human single-cell transcriptomic analyses of tumors revealed coordinated GABA_B_ and immune checkpoint signaling within a shared myeloid-centered communication network. Together, these findings suggest that inhibitory signaling contributes to the integration of neuronal and immune processes in GBM and may influence therapeutic responsiveness. However, the mechanistic relationship between these effects remains to be established.

Our findings extend current models of neuron–glioma interactions, which have largely focused on excitatory neuronal activity. Previous studies have established that GBM exploits excitatory neuronal signaling to promote tumor progression ^37,38^, whereas the contribution of inhibitory circuits remains less understood ^9,39^. It was recently found that glioma growth is accompanied by disruption of peritumoral inhibitory circuits, including loss of PV^+^ and SOM^+^ interneurons, reduced inhibitory markers, and impaired GABAergic neurotransmission ^10^.

Together with the present finding that activation of peritumoral PV^+^ interneurons delays glioma progression (**Fig.1**), these observations raise the possibility that disruption of local inhibitory circuits may contribute to the establishment of a neuronal environment permissive to tumor growth. Conversely, inhibition of PV^+^ interneurons primarily worsened neurological dysfunction without significantly affecting tumor progression, suggesting that these circuits may contribute not only to tumor control but also to the preservation of cortical function during disease progression (**Fig.2**).

Beyond their role in neuronal homeostasis, inhibitory circuits may participate in local neuroimmune regulation ^40^, and GABAergic signaling can directly influence immune-cell function. GABA has been shown to promote monocyte differentiation toward IL-10-producing immunoregulatory macrophages and to suppress antitumor T-cell responses ^13^. Our findings suggest that the consequences of GABAergic signaling in GBM may similarly depend on interactions among neuronal, tumor, and immune compartments. Consistent with this interpretation, baclofen delayed tumor progression in GL261-bearing mice but showed limited efficacy in the more aggressive CT-2A model (**Fig.3I-P**). These syngeneic models display distinct biological and immunological features, with CT-2A exhibiting a highly tumorigenic and relatively immunotherapy-resistant phenotype and GL261 displaying greater intrinsic immunogenicity and a more immune-reactive microenvironment ^41–44^. These differences may contribute to the model-dependent effects of baclofen and argue against a uniform antitumor activity of GABA_B_ receptor activation.

This context dependence is further supported by recent evidence showing sex-dependent protumoral effects of GABA_B_ signaling through granulocytic myeloid-derived suppressor cells (gMDSCs). Pathak et al. reported that GABBR agonism enhanced the T-cell-suppressive activity of gMDSCs and promoted tumor growth specifically in female mice, whereas GABBR antagonism reduced their immunosuppressive phenotype and prolonged survival ^14^. Although apparently divergent from our findings in GL261, these observations reinforce the concept that the biological consequences of GABA_B_ signaling are strongly context dependent. Tumor model, sex, treatment paradigm, and the composition or functional state of the immune microenvironment may all influence the response to GABAergic modulation.

Against this context-dependent and incomplete activity of baclofen monotherapy, the marked therapeutic response observed with combined baclofen and PD-L1 blockade was particularly evident. Indeed, combination therapy induced complete tumor regression in the majority of treated GL261-bearing mice, markedly prolonged survival, and established protection against tumor rechallenge (**Fig.4**). Early immunohistochemical analysis revealed changes in the phenotypic composition of intratumoral F4/80+ cells preceding radiological regression, with a reduced proportion of F4/80+Arg1+ cells and an increased proportion of F4/80^+^CD11c^+^ cells (**Fig.4J, L**). These changes may reflect a relative reduction in an Arg1-associated phenotype and enrichment of a CD11c-expressing state during therapeutic response. However, Arg1 and CD11c expression alone does not define the functional state of myeloid cells, and these observations should not be interpreted as evidence of functional polarization or a causal contribution to tumor regression. Thus, rather than supporting a simple model in which GABA_B_ activation is intrinsically antitumoral, our findings suggest that its therapeutic consequences emerge from interactions with the immune microenvironment and, in GL261, can markedly enhance responsiveness to PD-L1 blockade.

The human single-cell analyses provide independent support for a relationship between GABAergic and immune signaling within the GBM microenvironment. GABA_B_ and PD-L1 signaling co-varied across individual tumors along a shared axis dominated by macrophages, microglia, and monocytes, with highly weighted interactions including SPP1–CD44, APP– CD74, TREM2–TYROBP, and extracellular matrix-associated pathways (**Fig.5**). Although correlative, these analyses identify a myeloid-centered communication landscape in which GABAergic and immune checkpoint signaling converge. This observation complements the early macrophage-associated changes observed following combination therapy in mice, without implying direct pathway crosstalk or that identical cellular mechanisms operate in the two settings. The specific cellular populations through which baclofen mediates its therapeutic effects remain to be established.

These findings have potential translational implications. Baclofen is an FDA-approved GABA_B_ receptor agonist with an established safety profile and extensive clinical use. While our data do not support baclofen as an antitumor agent *per se*, they suggest that GABA_B_ receptor modulation can substantially enhance the efficacy of PD-L1 blockade in a responsive tumor context. At the same time, the model dependence observed here, together with the reported sex-specific protumoral effects of GABA_B_ activation, emphasizes that clinical translation will require identification of the cellular, immunological, and potentially sex-dependent determinants of response. Rather than supporting indiscriminate GABA_B_ activation in GBM, our findings provide a rationale for defining the biological contexts in which GABAergic modulation may be exploited to enhance immunotherapy.

However, several limitations should be considered. Our immunohistochemical analysis identifies early phenotypic changes in F4/80^+^ myeloid populations but does not establish their functional contribution to therapeutic response. Similarly, ligand–receptor analyses infer potential cell–cell communication from transcriptomic data and cannot establish direct mechanistic interactions. Future studies combining higher-dimensional immune profiling with cell-specific perturbation will be required to identify the cellular populations mediating the interaction between GABA_B_ activation and PD-L1 blockade. The mechanisms underlying the divergent responses of GL261 and CT-2A tumors, as well as the contribution of sex, across different treatment contexts, warrant further investigation.

In conclusion, our work identifies inhibitory neuronal signaling as a component of the GBM neuroimmune system and demonstrates that pharmacological activation of GABA_B_ receptors can substantially enhance the therapeutic efficacy of PD-L1 blockade in a responsive preclinical context. Together, these findings extend current models of GBM by implicating inhibitory neuronal and GABAergic signaling in the neuroimmune landscape beyond direct neuron–tumor interactions and provide a framework for defining when GABAergic modulation may be therapeutically exploited to enhance immunotherapy.

## Acknowledgements

The authors would like to thank Elena Novelli for help in acquiring immunofluorescence images, Francesca Biondi and Sara Ciampi for animal care. We acknowledge the Multi Modal Molecular Imaging (MMMI) Italian Node of the Eurobioimaging Research Infrastructure for supporting the 7 T high-field MRI facility at the “G.Monasterio” Foundation. Finally, we dedicate this work to the memory of Prof. Matteo Caleo, whose scientific guidance, mentorship, and inspiration profoundly shaped our research and continue to influence our work.

## Author Contributions

All authors contributed to the study conception and design. Material preparation, data collection and analysis were performed by Marta Scalera, Elisa De Santis, Filippo Rossi, Nicolò Meneghetti and Alessandra Flori. The first draft of the manuscript was written by Marta Scalera and all authors commented on previous versions of the manuscript. All authors read and approved the final manuscript.

## Data Availability

The datasets generated during and/or analysed during the current study are available from the corresponding author on reasonable request.

## Ethics approval

All animal experiments were conducted in accordance with the ARRIVE guidelines and the European Communities Council Directive 86/609/EEC and were approved by the Italian Ministry of Health (authorization nos. 981/2020-PR and 433/2026-PR).

## References

1. Ostrom QT, Price M, Neff C, et al. CBTRUS Statistical Report: Primary Brain and Other Central Nervous System Tumors Diagnosed in the United States in 2016-2020. Neuro Oncol. 2023;25(12 Suppl 2):IV1–IV99. doi:10.1093/NEUONC/NOAD149

2. Jayaram MA, Phillips JJ. Role of the Microenvironment in Glioma Pathogenesis. Annu Rev Pathol. 2024;19:181–201. doi:10.1146/ANNUREV-PATHMECHDIS-051122-110348

3. Venkatesh HS, Johung TB, Caretti V, et al. Neuronal activity promotes glioma growth through neuroligin-3 secretion. Cell. 2015;161(4):803–816. doi:10.1016/j.cell.2015.04.012

4. Wirsching HG, Weller M. Does Neuronal Activity Promote Glioma Progression? Trends in Cancer. 2020;6(1):1–3. doi:10.1016/j.trecan.2019.11.002

5. Ramachandran R, Jeans AF. Breaking Down Glioma-Microenvironment Crosstalk. Neurosci. 2024;31(2):177. doi:10.1177/10738584241259773

6. Hijazi S, Smit AB, van Kesteren RE. Fast-spiking parvalbumin-positive interneurons in brain physiology and Alzheimer’s disease. Mol Psychiatry. 2023;28(12):4954–4967. doi:10.1038/S41380-023-02168-Y

7. Meyer J, Yu K, Luna-Figueroa E, Deneen B, Noebels J. Glioblastoma disrupts cortical network activity at multiple spatial and temporal scales. Nat Commun 2024 151. 2024;15(1):4503-. doi:10.1038/s41467-024-48757-5

8. Gill BJA, Khan FA, Goldberg AR, et al. Single unit analysis and wide-field imaging reveal alterations in excitatory and inhibitory neurons in glioma. Brain. 2022;145(10):3666–3680. doi:10.1093/BRAIN/AWAC168

9. Tantillo E, Vannini E, Cerri C, et al. Differential roles of pyramidal and fast-spiking, GABAergic neurons in the control of glioma cell proliferation. Neurobiol Dis. 2020;141:104942. doi:10.1016/j.nbd.2020.104942

10. Spalletti C, Scalera M, Mori E, et al. Inhibitory circuit dysfunction as a potential contributor to cortical reorganization in Glioblastoma progression. Neurobiol Dis. 2025;213. doi:10.1016/J.NBD.2025.106997

11. Pombo Antunes AR, Scheyltjens I, Lodi F, et al. Single-cell profiling of myeloid cells in glioblastoma across species and disease stage reveals macrophage competition and specialization. Nat Neurosci. 2021;24(4):595–610. doi:10.1038/S41593-020-00789-Y

12. Miller TE, El Farran CA, Couturier CP, et al. Programs, origins and immunomodulatory functions of myeloid cells in glioma. Nat 2025 6408060. 2025;640(8060):1072–1082. doi:10.1038/s41586-025-08633-8

13. Zhang B, Vogelzang A, Miyajima M, et al. B cell-derived GABA elicits IL-10+ macrophages to limit anti-tumour immunity. Nat 2021 5997885. 2021;599(7885):471–476. doi:10.1038/s41586-021-04082-1

14. Pathak A, Sravya P, Colon B, et al. GABA signaling activation drives glioblastoma progression in female mice through myeloid-derived suppressor cells. Nat Cancer 2026 77. 2026;7(7):1080–1093. doi:10.1038/s43018-026-01192-5

15. Garofalo S, D’Alessandro G, Chece G, et al. Enriched environment reduces glioma growth through immune and non-immune mechanisms in mice. Nat Commun. 2015;6:6623. doi:10.1038/ncomms7623

16. Brooks SP, Dunnett SB. Tests to assess motor phenotype in mice: a user’s guide. Nat Rev Neurosci. 2009;10(7):519–529. doi:10.1038/NRN2652

17. Spalletti C, Alia C, Lai S, et al. Combining robotic training and inactivation of the healthy hemisphere restores pre-stroke motor patterns in mice. Elife. 2017;6:1–31. doi:10.7554/eLife.28662

18. Velíšková J, Velíšek L. Behavioral Characterization and Scoring of Seizures in Rodents. Model Seizures Epilepsy Second Ed. Published online January 1, 2017:111–123. doi:10.1016/B978-0-12-804066-9.00009-2

19. Pizzorusso T, Medini P, Berardi N, Chierzi S, Fawcett JW, Maffei L. Reactivation of ocular dominance plasticity in the adult visual cortex. Science (80-). 2002;298(5596):1248–1251. doi:10.1126/science.1072699

20. Porciatti V, Pizzorusso T, Maffei L. The visual physiology of the wild type mouse determined with pattern VEPs. Vision Res. 1999;39(18):3071–3081. doi:10.1016/S0042-6989(99)00022-X

21. Restani L, Cerri C, Pietrasanta M, Gianfranceschi L, Maffei L, Caleo M. Functional Masking of Deprived Eye Responses by Callosal Input during Ocular Dominance Plasticity. Neuron. 2009;64(5):707–718. doi:10.1016/j.neuron.2009.10.019

22. Vannini E, Olimpico F, Middei S, et al. Electrophysiology of glioma: a Rho GTPase-activating protein reduces tumor growth and spares neuron structure and function. Neuro Oncol. 2016;18(12):1634–1643. doi:10.1093/neuonc/now114

23. Ruiz-Moreno C, Salas SM, Samuelsson E, et al. Harmonized single-cell landscape, intercellular crosstalk and tumor architecture of glioblastoma. bioRxiv. Published online August 27, 2022:2022.08.27.505439. doi:10.1101/2022.08.27.505439

24. Gayoso A, Lopez R, Xing G, et al. A Python library for probabilistic analysis of single-cell omics data. Nat Biotechnol 2022 402. 2022;40(2):163–166. doi:10.1038/s41587-021-01206-w

25. Troulé K, Petryszak R, Cakir B, et al. CellPhoneDB v5: inferring cell–cell communication from single-cell multiomics data. Nat Protoc 2025 2012. 2025;20(12):3412–3440. doi:10.1038/s41596-024-01137-1

26. Dimitrov D, Schäfer PSL, Farr E, et al. LIANA+ provides an all-in-one framework for cell–cell communication inference. Nat Cell Biol 2024 269. 2024;26(9):1613–1622. doi:10.1038/s41556-024-01469-w

27. Fard N, Amir L, Chakit A, et al. Communication Breakdown and Evolution of the Cancer Cell. bioRxiv. Published online October 6, 2025:2025.08.15.670478. doi:10.1101/2025.08.15.670478

28. Jin S, Plikus M V., Nie Q. CellChat for systematic analysis of cell–cell communication from single-cell transcriptomics. Nat Protoc 2024 201. 2024;20(1):180–219. doi:10.1038/s41596-024-01045-4

29. Arora C, Matic M, Bisceglia L, et al. The landscape of cancer-rewired GPCR signaling axes. Cell Genomics. 2024;4(5):100557. doi:10.1016/j.xgen.2024.100557

30. Vannini E, Maltese F, Olimpico F, et al. Progression of motor deficits in glioma-bearing mice: impact of CNF1 therapy at symptomatic stages. Oncotarget. 2017;8(14):23539–23550. doi:10.18632/oncotarget.15328

31. Romito JW, Turner ER, Rosener JA, et al. Baclofen therapeutics, toxicity, and withdrawal: A narrative review. SAGE open Med. 2021;9. doi:10.1177/20503121211022197

32. Khasraw M, Reardon DA, Weller M, Sampson JH. PD-1 inhibitors: Do they have a future in the treatment of glioblastoma? Clin Cancer Res. 2020;26(20):5287. doi:10.1158/1078-0432.CCR-20-1135

33. Robert C, Schachter J, Long G V., et al. Pembrolizumab versus Ipilimumab in Advanced Melanoma. N Engl J Med. 2015;372(26):2521–2532. doi:10.1056/NEJMOA1503093;JOURNAL:JOURNAL:NEJMS;ISSUE:ISSUE:DOI

34. Reck M, Rodríguez-Abreu D, Robinson AG, et al. Pembrolizumab versus Chemotherapy for PD-L1-Positive Non-Small-Cell Lung Cancer. N Engl J Med. 2016;375(19):1823–1833. doi:10.1056/NEJMOA1606774

35. Nejo T, Krishna S, Yamamichi A, et al. Glioma-neuronal circuit remodeling induces regional immunosuppression. Nat Commun 2025 161. 2025;16(1):4770-. doi:10.1038/s41467-025-60074-z

36. Tomaszewski WH, Waibl-Polania J, Chakraborty M, et al. Neuronal CaMKK2 promotes immunosuppression and checkpoint blockade resistance in glioblastoma. Nat Commun 2022 131. 2022;13(1):6483-. doi:10.1038/s41467-022-34175-y

37. Venkataramani V, Tanev DI, Strahle C, et al. Glutamatergic synaptic input to glioma cells drives brain tumour progression. Nature. 2019;573(7775):532–538. doi:10.1038/s41586-019-1564-x

38. Venkatesh HS, Morishita W, Geraghty AC, et al. Electrical and synaptic integration of glioma into neural circuits. Nature. 2019;573(7775):539–545. doi:10.1038/s41586-019-1563-y

39. Blanchart A, Fernando R, Häring M, et al. Endogenous GAB AA receptor activity suppresses glioma growth. Oncogene. 2017;36(6):777–786. doi:10.1038/onc.2016.245

40. Piletti Chatain C, Gursky ZH, Camacho DF, Favuzzi E. Immunity meets inhibition: Immunomodulation of cortical inhibitory synaptic networks. Curr Opin Neurobiol. 2025;94. doi:10.1016/J.CONB.2025.103090

41. Mikolajewicz N, Tatari N, Wei J, et al. Functional profiling of murine glioma models highlights targetable immune evasion phenotypes. Acta Neuropathol. 2024;148(1). doi:10.1007/S00401-024-02831-W

42. Yadav N, Purow BW. Understanding current experimental models of glioblastoma-brain microenvironment interactions. J Neurooncol. 2024;166(2):213–229. doi:10.1007/S11060-023-04536-8

43. Iorgulescu JB, Ruthen N, Ahn R, et al. Antigen presentation deficiency, mesenchymal differentiation, and resistance to immunotherapy in the murine syngeneic CT2A tumor model. Front Immunol. 2023;14. doi:10.3389/FIMMU.2023.1297932

44. Khalsa JK, Cheng N, Keegan J, et al. Immune phenotyping of diverse syngeneic murine brain tumors identifies immunologically distinct types. Nat Commun 2020 111. 2020;11(1):1–14. doi:10.1038/s41467-020-17704-5

