## Supplementary Material for "GABAergic Circuit Activation Induces a Therapeutically Responsive State for anti-PD-L1 Immunotherapy in Glioblastoma"

**Supplementary Data**

**Table 1**

| Gene | Primer | Sequence | Tm (°C) | Product size (bp) |
|---|---|---|---|---|
| *Actb* | Forward | 5’ TACCACCATGTACCCAGGCATT 3’ | 60 | 188 |
| | Reverse | 5’ ACTCATCGTACTCCTGCTTGCTGA 3’ | 60 | |
| *Gabbr1* | Forward | 5’ ACAGACCAAATCTACCGGGC 3’ | 63.1 | 152 |
| | Reverse | 5’ GTGCTGTCGTAGTAGCCGAT 3’ | 63.1 | |
| *Gabbr2* | Forward | 5’ AAGCTCAAGGGGAACGACG 3’ | 63.1 | 106 |
| | Reverse | 5’ ACTTGCTGCCAAACATGCTC 3’ | 63.1 | |

**Table 1. Primer sequences used for real-time PCR analysis.** Forward and reverse primer sequences for Actb, Gabbr1, and Gabbr2 are shown in the 5′→3′ direction, together with their melting temperatures (Tm, °C) and expected amplicon sizes (bp).

### Supplementary figure 1

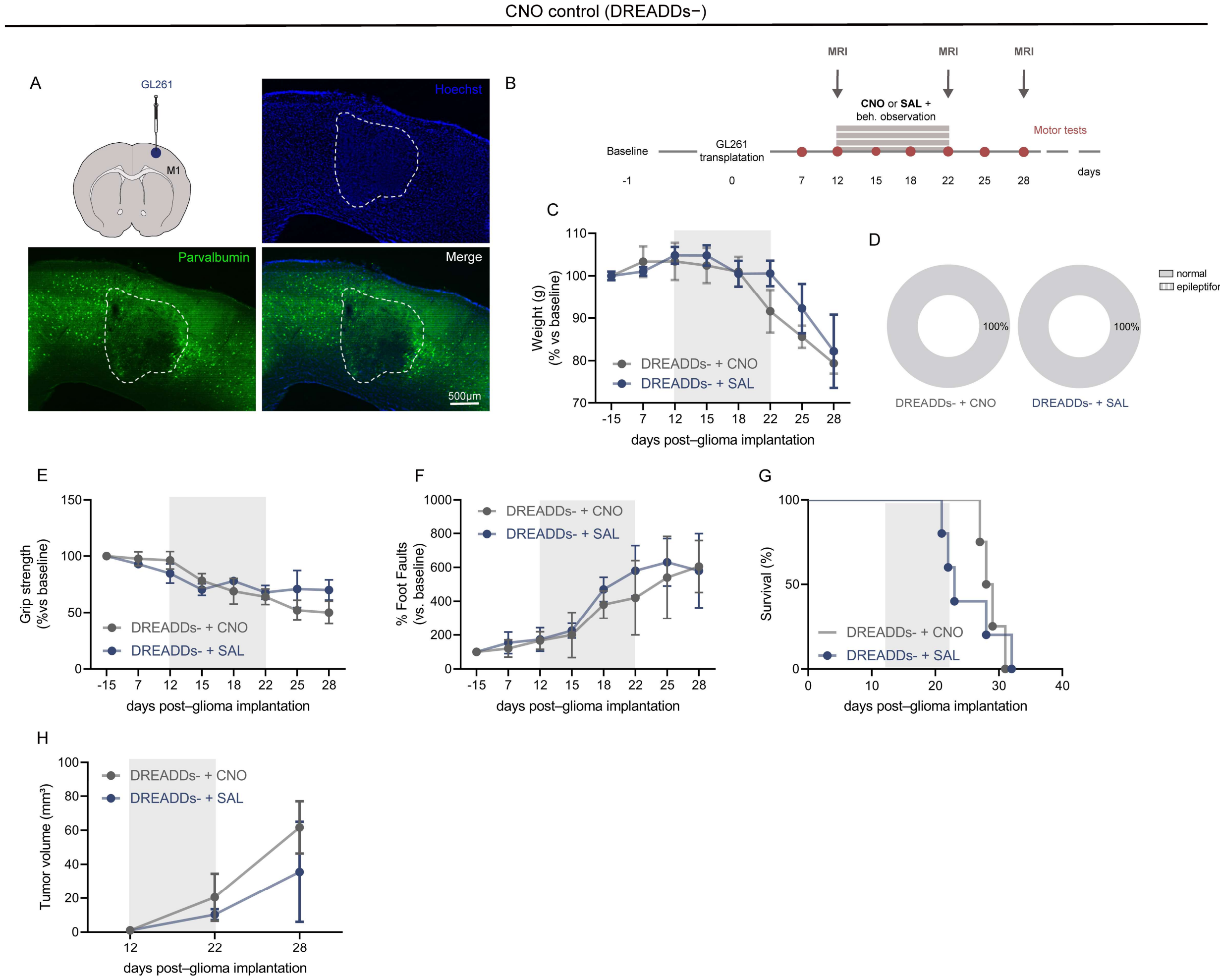


**Supplementary Figure 1. Chronic CNO administration in the absence of DREADD expression does not affect tumor progression, animal physiology, or motor performance.** (**A**) Schematic of GL261 glioma cell transplantation into the cortex of mice that did not receive adeno-associated virus (AAV) injections for DREADD expression (left), and representative immunofluorescence brain coronal sections showing Hoechst nuclear staining (blue, top right), Parvalbumin-positive interneurons (green, bottom left), and merged channels (bottom right) delineating the tumor area (dashed line). Scale bar = 500 µm. (**B**) Experimental plan. CNO = clozapine N-oxide (1 mg/kg, i.p.); SAL = saline (vehicle). (**C**) Longitudinal monitoring of body weight changes, expressed as a percentage of baseline values (day -15), indicating stable and comparable physiological conditions for CNO-treated mice (DREADDs– + CNO, grey line, n = 5) and vehicle controls (DREADDs– + SAL, blue line, n = 5). (**D**) Donut charts illustrating behavioral observation outcomes, with 100% normal clinical conditions in both the DREADDs– + CNO (grey chart) and DREADDs– + SAL (blue chart) groups. (**E**) Grip strength and (**F**) grid walk (% foot faults) performance across tumor progression, demonstrating a comparable motor decline in both cohorts. Data in C, E, and F represent means ± SEM, analyzed by two-way repeated-measures ANOVA, normalized to baseline values. (**G**) Kaplan–Meier survival curves showing no significant difference in overall survival between the DREADDs– + CNO (grey line, n = 5) and DREADDs– + SAL (blue line, n = 5) groups (log-rank Mantel–Cox test). (**H**) Volumetric quantification of longitudinal tumor growth via MRI, demonstrating identical kinetics of tumor expansion between the two groups. Values are normalized to day 12 (two-way repeated-measures ANOVA).

**Supplementary figure 2**

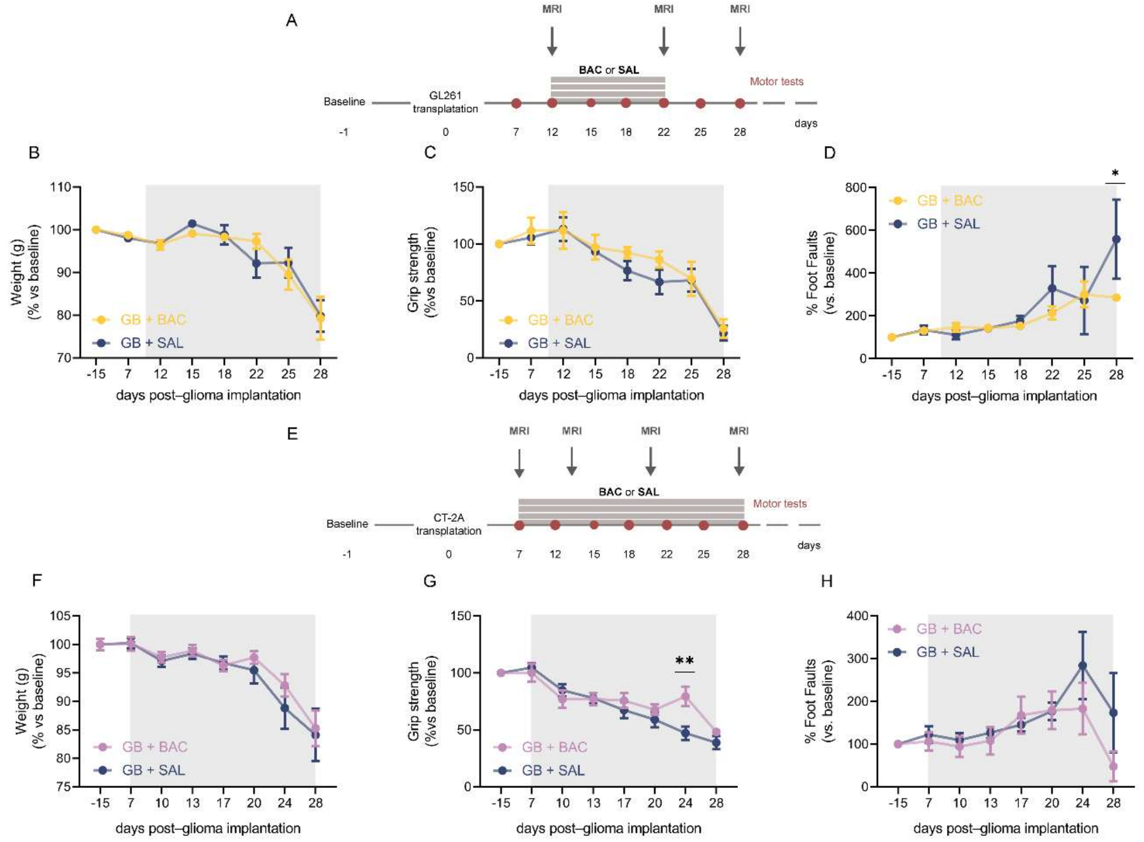


**Supplementary Figure 2. Impact of baclofen monotherapy on physiological parameters and motor performance in GL261 and CT-2A glioma models.** (**A**) Experimental timeline illustrating the paradigm for GL261 glioma-bearing mice subjected to longitudinal MRI, vehicle (SAL), or baclofen (BAC) pharmacotherapy, and motor assessment. BAC = baclofen (1 mg/kg, i.p.); SAL = saline (vehicle). (**B**) Longitudinal monitoring of body weight changes, expressed as a percentage of baseline values (day -15), indicating comparable physiological decline between baclofen-treated mice (GB + BAC, yellow line, n = 8) and vehicle controls (GB + SAL, blue line, n = 6) in the GL261 model. (**C**) Grip strength and (**D**) grid walk (% foot faults) performance across GL261 tumor progression, demonstrating a significant reduction in foot faults at the final experimental timepoint in the baclofen cohort compared with the vehicle group (D; $* P < .05$ at day 28). (**E**) Schematic representation of the in vivo experimental design and treatment schedule optimized for the CT-2A syngeneic model. (**F**) Longitudinal body weight changes in CT-2A glioma-bearing mice, revealing no significant differences between the GB + BAC (violet line, n = 8) and GB + SAL (blue line, n = 8) cohorts. (**G**) Grip strength and (**H**) grid walk performance in the CT-2A model, showing a transient preservation of grip strength in baclofen-treated mice at advanced stages of tumor progression (G; $P < .01$ at day 24). Data in B–D and F–H represent means ± SEM, analyzed by two-way repeated-measures ANOVA, normalized to baseline values.

### Supplementary figure 3

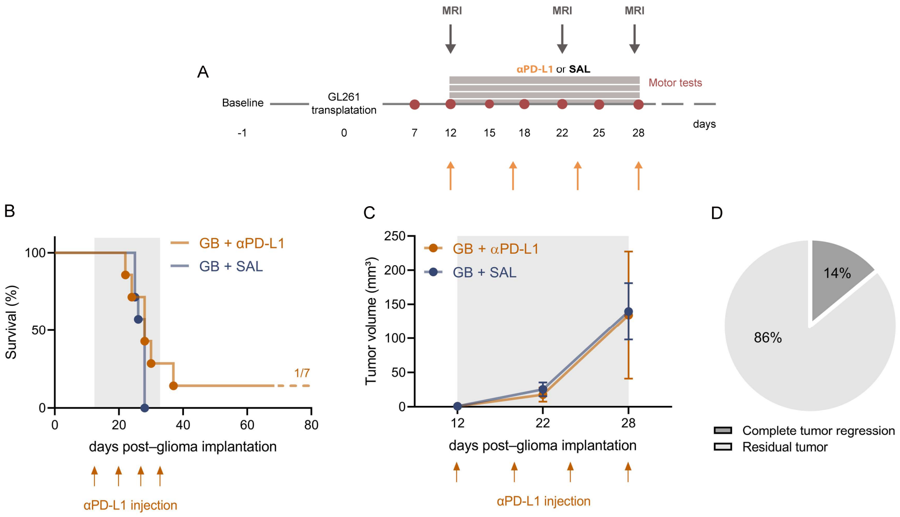


**Supplementary Figure 3. Evaluation of anti-PD-L1 monotherapy in the GL261 glioma model.** (**A**) Experimental timeline schematic illustrating GL261 glioma cell transplantation at day 0. Mice in these control cohorts received either anti-PD-L1 antibody (αPD-L1) monotherapy or saline (SAL) vehicle control twice weekly from day 12 to day 28 post-implantation (orange arrows), in the absence of systemic baclofen. Arrows indicate time points for magnetic resonance imaging (MRI) acquisition and behavioral motor testing. (**B**) Kaplan-Meier survival analysis of GL261-bearing mice. Anti-PD-L1 monotherapy (αPD-L1, orange line) yielded a long-term survival rate of only 14% (n = 1/7). Vehicle controls (SAL, blue line) exhibited 100% mortality. The grey shaded area defines the treatment duration. (**C**) Longitudinal quantification of tumor volume ($mm^3$) determined by MRI at days 12, 22, and 28 post-transplantation. Both the αPD-L1 monotherapy (orange) and SAL vehicle (blue) groups showed progressive tumor burden, contrasting with the regression observed under combination therapy. Data represent mean ± SEM. (**D**) Pie chart quantifying the best radiological response within the anti-PD-L1 monotherapy cohort, demonstrating complete tumor regression in 14% (dark grey) and persistent residual tumor in 86% (light grey) of analyzed animals.
